# A protein-centric approach for exome variant aggregation enables sensitive association analysis with clinical outcomes

**DOI:** 10.1101/653683

**Authors:** Ginny X.L. Li, Dan Munro, Damian Fermin, Christine Vogel, Hyungwon Choi

**Author notes:** Corresponding author: Hyungwon Choi.

## Abstract

Somatic mutations are early drivers of tumorigenesis and tumor progression. However, the mutations typically occur at variable positions across different individuals, resulting in the data being too sparse to test meaningful associations between variants and phenotypes. To overcome this challenge, we devised a novel approach called Gene-to-Protein-to-Disease (GPD) which accumulates variants into new sequence units as the degree of genetic assault on structural or functional units of each protein. The variant frequencies in the sequence units were highly reproducible between two large cancer cohorts. Survival analysis identified 247 sequence units in which somatic mutations had deleterious effects on overall survival, including consensus driver mutations obtained from multiple calling algorithms. By contrast, around 75% of the survival predictive units had been undetected by conventional gene-level analysis. We demonstrate the ability of these signatures to separate patient groups according to overall survival, therefore providing novel prognostic tools for various cancers. GPD also identified sequence units with somatic mutations whose impact on survival was modified by the occupancy of germline variants in the surrounding regions. The findings indicate that a patient’s genetic predisposition interacts with the effect of somatic mutations on survival outcome in some cancers.

## Introduction

Accumulation of mutations in the genome can impair cellular function, lead to abnormal proliferation of cells, and eventually cause cancer (Martincorena and Campbell, 2015). Today’s massively parallel sequencing technology can detect such variation in the whole genome or exome, and simultaneous analysis of paired tumor and normal samples enables reliable calling of somatic mutations and germline variants. Large-scale sequencing projects such as The Cancer Genome Atlas (TCGA) have successfully mapped the mutation landscape of >11,000 patients across 33 cancer types (Ding et al., 2018; Ellrott et al., 2018; Huang et al., 2018; Liu et al., 2018). The next logical step is to test for the association between cancer genome variants and clinical outcomes such as prognosis and response to first-line therapies (Castro et al., 2013; Gentles et al., 2015; Dellinger et al., 2016). For example, somatic driver mutations are responsible for tumor maintenance, progression, and metastasis (Reimand and Bader, 2013), which are robust indicators of patient survival.

Indeed, mutations at specific loci can make a marked difference in disease prognosis. For example, tumors from papillary thyroid carcinoma patients with the missense mutation V600E in the *BRAF* gene are often associated with more rapid tumor growth and higher death risk than the patients without the mutation (Xing, 2005), Further, missense mutations affect the transcriptional activity of tumor suppressor TP53 in different ways, as has been shown in a study of ~1,800 primary breast cancer patients (Olivier, 2006). Mutations found within the DNA-binding motif, especially those at codon 179 and 248, are associated with poorer patient survival outcome than mutations at other positions. Other than these well-known examples, however, systematic identification of variants useful for prognosis still remains a major undertaking of cancer genomics.

In the past, a plethora of algorithms have been developed to differentiate truly cancer causing, driver mutations from abundant passenger mutations. MuSic (Dees et al., 2012), OncodriveCLUST (Tamborero et al., 2013), and MutSigCV (Lawrence et al., 2013) identify the genes that harbor significantly more mutations than others based on cancer-specific background mutations; SIFT (Kumar et al., 2009), FATHMM (Shihab et al., 2013), PolyPhen2 (Adzhubei et al., 2010), and CHASM (Wong et al., 2011) identify mutations by evaluating their functional impact; e-Driver (Porta-Pardo and Godzik, 2014) and ActiveDriver (Reimand and Bader, 2013) focus on mutations residing at structurally important sites; and Network-Based Stratification (Hofree et al., 2013), PARADIGM (Vaske et al., 2010), TieDIE (Paull et al., 2013), and DriverNet (Bashashati et al., 2012) prioritize driver mutations using network- and pathway-based approaches. In a recent massive meta-analysis of the Pan-Cancer Atlas project, Bailey *et al. (Bailey et al., 2018)* applied 26 different computational tools and identified 299 consensus driver genes and ~3,400 mutations across 33 cancer types -- one of the most comprehensive such studies known to date.

Despite these developments, it remains challenging to test statistical association between mutations and meaningful clinical outcomes such as patient prognosis. One reason behind this challenge is the extreme heterogeneity of tumor genomes (McGranahan and Swanton, 2015). In contrast to studies of single-nucleotide polymorphisms in healthy individuals, somatic mutations detected in tumor specimens rarely reside at the same locus across many patients (Pickrell et al., 2016). As a result, most tumor variants have very low frequency, e.g. on average <0.011% across TCGA Pan-Cancer Atlas patients, with most of them detected in a single patient. Further, the issue potentially reflects the varying genome instability across patients, as is typical for cancer cells (Jeggo et al., 2016). As a consequence, statistical analysis of somatic mutations at the locus level inherently lacks the statistical power to detect prognostic markers (Kiiski et al., 2016).

A common remedy to this challenge is to count mutations *per* gene, or to classify the mutation status of a gene as ‘mutated’ or ‘not mutated’ in each patient (Candido-dos-Reis et al., 2015). A disadvantage of this approach is that it considers patients with different mutation profiles as carriers of the same mutation burden, regardless of the positions of mutations. An alternative systems-oriented approach, often referred to as network-based stratification methods, propagates the binary mutation status of genes onto a protein-protein interaction network and identifies interacting genes enriched with adjacent mutation events as etiologically important players (Hofree et al., 2013; Zhong et al., 2015; Hristov and Singh, 2017; Le Morvan et al., 2017). While this method more sensitively identifies mutation clusters than locus-based analysis, it ignores the relative positions of mutations in the gene sequence and subsequently fails to provide directly testable hypotheses on the gain or loss of protein function.

To counter these challenges and enable sensitive identification of prognostic exome variants and preserve information on the location of the mutations, we developed a new approach called Gene-to-Protein-to-Disease (GPD), which maps somatic mutations (and/or germline variants) onto small sequence units located in protein coding or non-coding regions. The approach therefore transforms the original variant data into substantially less sparse count data and enables statistical analysis for disease prognosis. It successfully recovers a handful of cancer driver mutations as prognostic markers, and also enables identification of interactions between the effects of somatic mutations and germline variants on survival outcomes.

## Materials and Methods

### Data Source

Exome sequencing data for somatic mutations in TCGA datasets were downloaded from the Genomic Data Commons (MC3 Public MAF) (Ellrott et al., 2018). Germline variant data were downloaded from the ISB-CGC web interface, where the data is deposited in a VCF file format (Huang et al., 2018). We further annotated the file with the snpEff tool (genome version GRch37.75) (Cingolani et al., 2012). In addition, we gathered 348,658 protein modification sites (244,852 phosphorylation, 46,352 ubiquitination, 25,807 acetylation, 20,691 methylation, 8,388 SUMOylation, and 2,568 glycosylation sites) from PhosphoSitePlus (Hornbeck et al., 2015), and 45,607 domains, families and repeats for 19,076 genes from Pfam (El-Gebali et al., 2019).

Survival outcome data were downloaded from TCGA Pan-Cancer Clinical Data Resource (Liu et al., 2018), which contains curated clinical information for 10,793 patients. Among these, 367 non-primary skin cutaneous melanoma patients with metastatic tumors were excluded. The data resource provides four types of clinical endpoints, including overall and disease-specific survival (OS, DSS), as well as disease-free and progression-free time interval (DSI, PFI). As advised by Liu et al., OS and PFI are the most reliable endpoints for the majority of the cancers. Hence, we report the results for the OS as the endpoint of interest in this study, while reporting the results for PFI in the **Supplementary Information**. Abbreviations for cancer types have been adapted from TCGA available at https://gdc.cancer.gov/resource-tcga-users/tcga-code-table/tcga-study-abbreviations.

### Mapping scheme

GPD segments exons and adjacent genomic regions of each gene into three types of units (**Figure 1A**). First, Protein Information Units (PIU) refer to the genomic regions encoding protein domains, e.g., as defined by the Pfam database (El-Gebali et al., 2019)) or +/−5 amino acid-long windows around protein modification sites (e.g., as reported in the PhosphoSitePlus database). Sequence regions between PIUs are defined as linker units (LU). The LUs not only include linker regions between domains, but also cover unannotated, repeat or disordered regions. The regions outside the protein-coding sequences, including untranslated regions, introns, and regulatory regions are collectively defined as non-coding units (NCU). NCUs are assigned to the closest gene in the genome.

**Figure 1.**
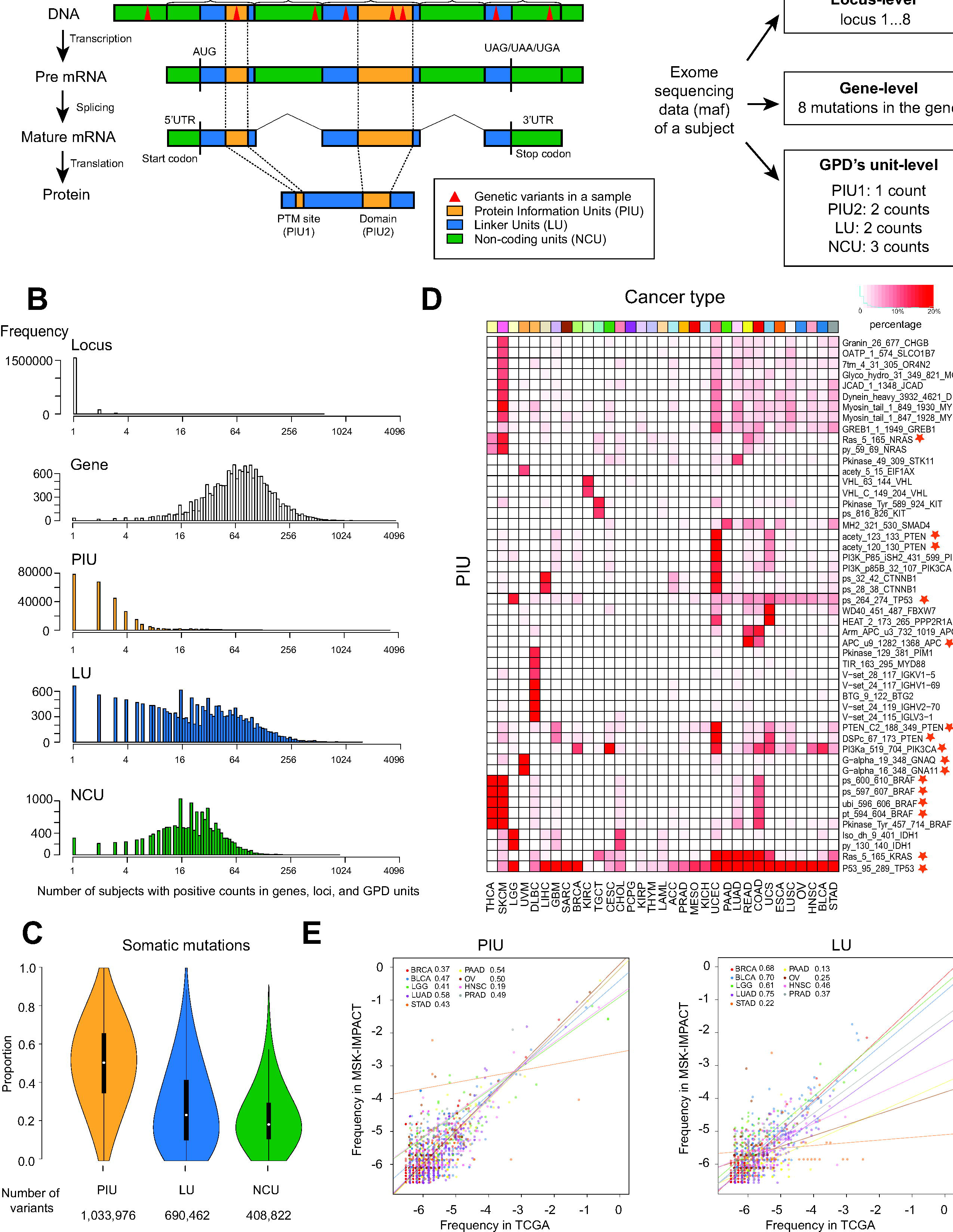
GPD mapping scheme and mapping results of somatic mutations across 33 cancers. (**A**) GPD’s protein information units (PIUs) consist of domains and windows surrounding protein modification sites (+/− 5 amino acids around a PTM site) in proteins (yellow); Regions of the protein sequence that are not covered by a domain and a PTM site-containing window are called linker units (LUs) (blue). All sequence regions that do not encode the final protein product are named non-protein coding units (NCUs). Mutations (red triangles) are mapped to the GPD units defined above. GPD’s unit-level data are per unit counts of mutations. Locus-level and gene-level counts are also illustrated for comparison. (**B**) Histograms of the mutation-carrying loci, genes, PIUs, LUs and NCUs in 10,793 subjects (x-axis) across all TCGA cancers. For example, a majority of mutations occurring at a specific locus have frequency of one or two individuals. By contrast, genes with any somatic mutations have a median count of 64. (**C**) Percentage of mutations mapped to one of the three types of sequences units across all genes. (**D**) Percentage of patients carrying mutations in the 50 most frequently mapped PIUs across 33 cancers. Stars indicate the PIUs mentioned in the main text. (**E**) Mutation frequency (logarithm base 2) of each PIU and LU among patients in TCGA Pan-Cancer cohort and MSK-IMPACT cohort.

Synonymous variants which do not result in amino acid changes in protein sequences were excluded from the analysis. Missense mutations resulting in single amino acid changes represent 68.2% of all genetic alterations in the data. These missense mutations are unambiguously mapped to sequence units and counted. For truncating mutations or those that can introduce large sequence changes (e.g. unexpected stop codon), GPD counts them the same way as it counts missense mutations, since it is difficult to quantify the effect of these mutations in terms of the mutational load on sequence units.

To understand the proximity between non-coding variants and DNA coding regions, we downloaded a GTF file on GRCh37 human genome annotation from GENCODE website (https://www.gencodegenes.org/human/release_32lift37.html). From this GTF file, we extracted the coordinates of start and stop codons for each gene. For non-coding variants outside of the translation interval, we calculated the distance (in base pairs) between the variant loci and the first base of start codons if they lie in the upstream region of the translation interval or the last base of stop codons if they are in the downstream region of the translation interval.

### Validation of mapping with MSK-IMPACT data

MSK-IMPACT cohort data was retrieved from https://www.cbioportal.org/study/summary?id=msk_impact_2017. This data includes exome-sequencing data from a total of 10 thousand patients across 14 cancer types. We implemented GPD mapping scheme to map mutations occurred in primary tumor samples collected in the study to PIUs, LUs and NCUs. Then we computed per sample mutation counts (mutation frequency) in each common PIU and LU mapped by both TCGA data and MSK-IMPACT data. Spearman correlations were calculated for each pair of mutation frequencies to measure the correlation between mapping results generated from two independent datasets.

### Selection of prognostic sequence units (main effect model)

The association between mutation count of each sequence unit and OS (or PFI) was evaluated using Cox proportional hazard regression. To focus on the association with long-term survival outcome, we excluded patients with less than 90 days survival time. For each cancer, we included sequence units with mutations in three or more patients for all subsequent analyses. We adjusted all regression models for the total mutation count, age, gender, race and tumor stage of each patient (available in ACC, BLCA, BRCA, CHOL, COAD, ESCA, HNSC, KICH, KIRC, KIRP, LIHC, LUAD, LUSC, MESO, PAAD, READ, SKCM, STAD, TGCT, THCA, UVM). The Cox model posits the hazard function as a product of the baseline risk function at time *t* and the relative risk based on covariates. For each sequence unit, the hazard function for subject *i* is given by:

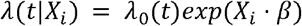

Where λ_0_ (*t*) denotes the baseline hazard function and *X*_*i*_ = {*Count*_*i*_, *TotalMutation*_*i*_, *Age*_*i*_, *Gender*_*i*_, *Race*_*i*_,*Stage*_*i*_} are the covariates. For comparison, we repeated the analysis but counted mutations *per* gene (gene-level analysis), instead of *per* sequence unit. For each model, the proportional hazard assumption was tested by evaluating the association between transformed survival time and the scaled Schoenfeld residuals (Schoenfeld, 1982).

### Validation of sequence unit signatures for the predictability of survival outcomes

We conducted 10-fold cross validation with supervised principal component method (Bair et al., 2006) for each cancer cohort to validate the robustness of predictability of the signatures. In each iteration of the 10-fold of validation, we used 9 partitions of randomly split samples as training data. We identified sequence units significantly associated with OS (*q*-value 0.05) and synthesized up to five principal components (PC) of the selected units and fit another cox proportional hazards model consisting of the covariates *X*_*i*_ = {*PC*1_*i*_, *PC*2_*i*_, *PC*3_*i*_, *PC*4_*i*_, *PC*5_*i*_, *TotalMutation*_*i*_, *Age*_*i*_, *Gender*_*i*_, *Race*_*i*_,*Stage*_*i*_}.

After fitting the models, we computed the predictive index (PI) for each training sample as a measurement of survival risk.

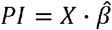

where 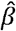 represents coefficients from the above models. Using the remaining one-fold of the data as test data, we first projected them onto the PCs with the loadings of principal components derived with training data, and computed the PIs for test samples as described above. Finally, test samples were clustered into the high-risk group if their PI is higher than the median of training samples’ PIs or the low-risk group otherwise. After iterating the process 10 times, we have assigned the samples into high- or low-risk groups. We then test the difference in survival between these two groups with log-rank test.

To investigate the degree of overfitting, we shuffled the survival outcomes among the samples 1,000 times. For each shuffled data, we conducted the 10-fold cross validation described above. This process generates 1,000 log-rank statistics, allowing us to form the empirical null distribution of the statistic, representing the assumption that there is no association between sequence unit-level mutation counts and survival outcomes. Therefore, the significance of survival difference in the original data can be derived by locating the log-rank statistic in the empirical distribution.

### Identification of interactions between effects

The hazard models accounting for the interaction between somatic mutations and germline variants consider three main covariates: the somatic mutation count of a unit, the total count of germline variants in the same gene, and the interaction between the two. Similar to the main effect models above, models were adjusted for total mutation count, gender, age, race and tumor stage. The hazard function is identical to the above, but denotes a vector of covariates as:

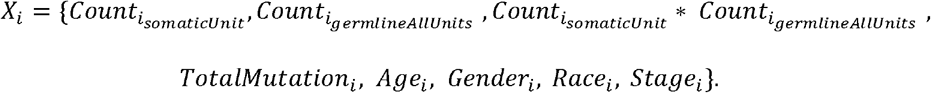

### GPD software implementation

The GPD framework was developed as an R package. The package is publicly available at https://github.com/ginnyintifa/GPD, with user-defined GPD units and control parameters for mapping and analysis. A user manual is available at the aforementioned GitHub site. The GPD implementation maps mutations to sequence units. The default genomic coordinates of protein domains and protein modification sites are provided in the package, but the user can also provide the information from other sources in the specified format. Users need to provide mutation data.

## Results

### GPD maps variants to “sequence units” encoding well-defined functional segments of protein sequences

**Figure 1A** illustrates the concept of GPD’s sequence units through a hypothetical exome with somatic mutations: two mutations occurred within a protein domain (PIU1), one mutation near a protein modification site (PIU2), two mutations in a LU outside the PIUs, and three mutations in the non-coding unit. Using the protein domains from Pfam (El-Gebali et al., 2019) database and the protein modification sites from PhosphoSitePlus (Hornbeck et al., 2015) database as PIUs, GPD mapped all somatic mutations extracted from TCGA to a total of 270,470 PIUs, 17,252 LUs, and 17,266 NCUs in 17,744 human genes, respectively.

Next, we counted the coding and non-coding variants in sequence units *per* patient, where the somatic mutations were called by TCGA Pan-Cancer Atlas (Ellrott et al., 2018). Mapping to GPD’s sequence units revealed that different cancers showed unique somatic mutation frequencies. For example, uterine cancer (UCEC) had on average >1,000 mutations *per* patient; skin and colon cancers (SKCM and COAD) had on average >500 mutations *per* patient. By contrast, the exomes in many other cancers (e.g. KICH, UVM, TGCT, THCA and PCPG) had fewer than 30 somatic mutations *per* patient on average.

As well known in the literature, the overall distribution of somatic mutations in the exomes of individuals was remarkably sparse and heterogeneous. Less than 8% of the somatic mutations appeared in two or more patients at the same locus (**Figure 1B**). As expected, the sparsity in the data due to the lack of common loci was significantly reduced when accumulating mutations *per* gene or *per* sequence unit. While 99% of the genes showed mutations, we found that 68% of PIUs, 96% of LUs, and 98% NCUs, respectively, harbored somatic mutations in two or more patients. The variable mutation frequencies across different sequence units within the same gene suggest that GPD’s mapping approach provides a higher-resolution map than gene-level mapping, as it can focus on smaller sequence regions while still preserving the coverage and statistical power to compare patient groups.

When examining the distribution of mutations across different genes, we noticed that almost half the somatic mutations were within PIUs (49%), one third were within LUs (32%), and 19% were in NCUs (**Figure 1C**). Among the mutations mapped to NCUs, 32% lie between start codon and stop codon, indicating that they are located in introns. For the non-intronic mutations in NCUs, 23% of the mutations are positioned upstream of a start codon and 77% are positioned downstream of a stop codon. A closer investigation of the distance between these non-intronic loci outside the coding regions to either the nearest start codon or stop codon revealed that 78% of them are located within 1,000 base pairs from either start codon or stop codon, and 92% are within 3,000 base pairs, implying that they are positioned in promoter regions, 5’ UTR, or 3’ UTR (**Supplementary Figure 1**). In the current definition of GPD’s sequence units, 67% of the protein coding sequence is covered by PIUs and 33% is covered by LUs. Accordingly, within the protein coding regions, both unit types have about equal chance to carry a mutation: the expected frequency of a mutation is 0.17 and 0.16 for PIUs and linker units, respectively.

**Figure 1D** shows the PIUs with the largest frequency of somatic mutations in TCGA patients. Not surprisingly, these PIUs are located in well-known cancer genes. For example, the PIU for the DNA-binding domain of *TP53* gene showed consistent somatic mutations across the majority of the cancers. Other mutations surrounded the serine phosphorylation site S269 of the protein, which is crucial for the conformational stability of the protein (Fraser et al., 2010). In another gene, *KRAS*, we found the highest frequency of somatic mutations in the PIU corresponding to the RAS domain of the protein in pancreas, lung, rectum, and colon cancer (PAAD, LUAD, READ, and COAD). The RAS domain also accumulates most mutations in the *NRAS* gene in 103 non-metastatic skin cancer (SKCM). Further, we find frequent mutations in thyroid, skin, and colon cancers (THCA, SKCM, and COAD) in a region in the *BRAF* gene densely populated with phosphorylation and ubiquitination sites (T599, K601, S602, S605, and K601). Similarly, the cancer gene *PTEN* accumulated mutations around lysine acetylation sites K125 and K128. **Supplementary Table 1** enumerates the sequence units with high mutation rates in specific cancers.

Next, to assess the reproducibility of the mapping scheme by sequence units, we validated the results above (see **Materials and Methods**) with somatic mutation data for nine cancers from an independent cohort (MSK-IMPACT) (Zehir et al., 2017). As mutations found outside protein coding regions (i.e. NCUs) were excluded in MSK-IMPACT, we mapped variants to PIUs and LUs only. We found that the somatic mutation frequencies in the sequence units are highly reproducible between the TCGA and MSK-IMPACT cohorts (**Figure 1E**), suggesting that somatic mutations have the propensity to be enriched in specific GPD units within each gene.

### Prognostic signatures of somatic mutations for overall survival are validated in eight cancer types

To identify sequence units harboring somatic mutations of prognostic potential, we tested each sequence unit for positive association with OS in Cox proportional hazard models (see **Materials and Methods**). We discovered 196, 170, and 107 unique PIUs, LUs and NCUs that were associated with OS in at least one of 24 cancer types, with mutation(s) occurring in at least three patients (*q*-value 0.05). Also, Schöenfeld residuals suggested no violation of proportional hazards assumption in these sequence units. We performed 10-fold cross validation on each cancer type (see **Materials and Methods**) to examine if the discovered prognostic units are capable of predicting survival outcomes in a reproducible manner. In eight cancer types (BLCA, BRCA, ESCA, GBM, KIRC, LGG, OV, UCEC), the predictive indices can cluster patients into two risk groups with significantly different survival outcomes (**Figure 2** and **Table 1**). Moreover, we conducted a permutation test to evaluate the degree of overfitting in the cross-validation process (see **Materials and Methods**). These eight cancers survived the test with no sign of overfitting (**Table 1** and **Supplementary Figure 2**). Therefore, our final prognostic signatures consist of 103 PIUs, 86 linker and 59 non-coding units discovered in these cancer cohorts (**Supplementary Table 2**).

**Figure 2.**
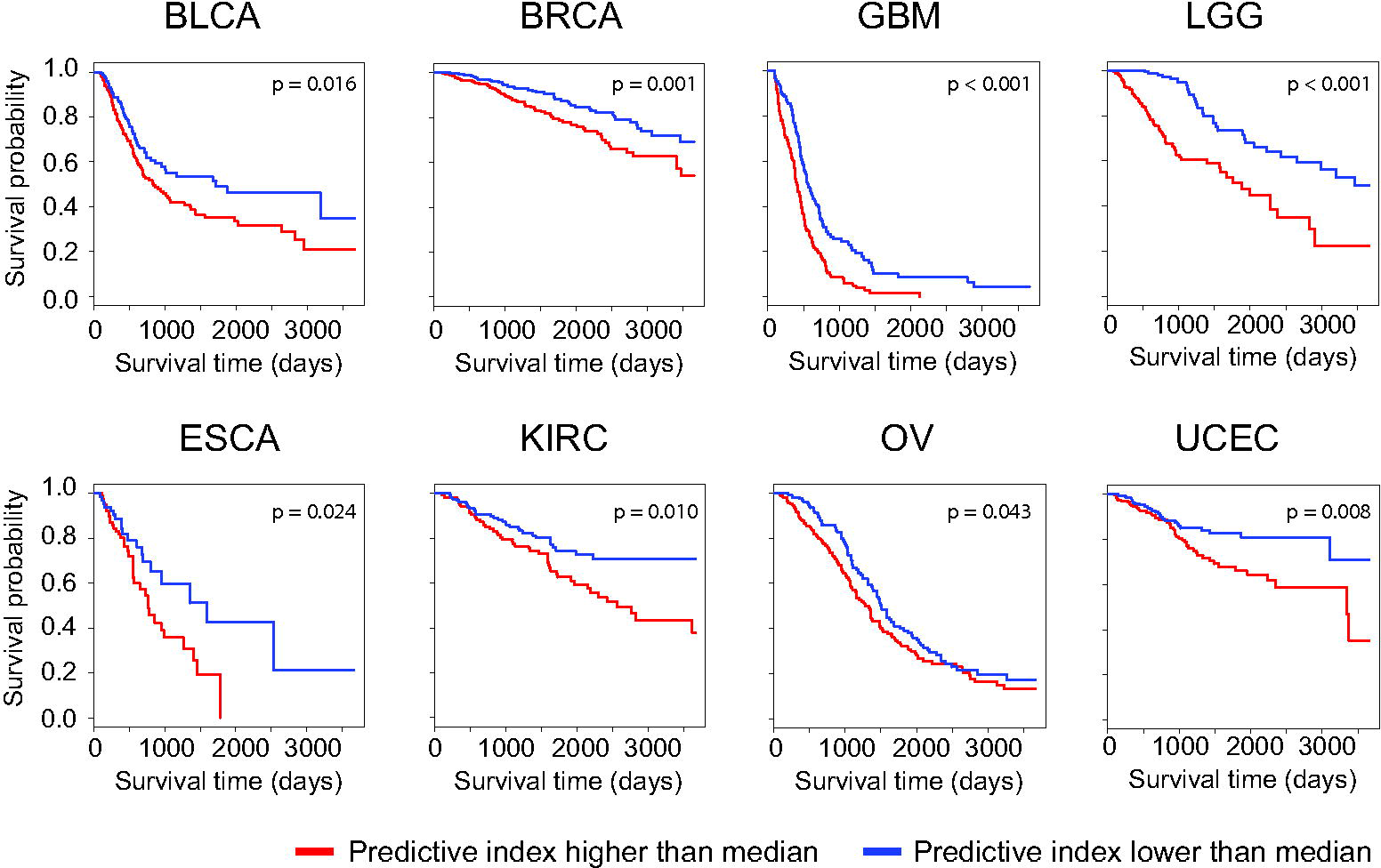
Kaplan-Meier plot for eight cancer types with validated prognostic sigantures of sequence units. Predictive index (PI) was derived for each patient in the test group in each fold of the 10-fold cross validation. Patients who have PIs greater than the median in training group are clustered in the high-risk group (red), those with lower than the median PI are in the low-risk group (blue). P-values are derived from log-rank tests.

**Table 1.**
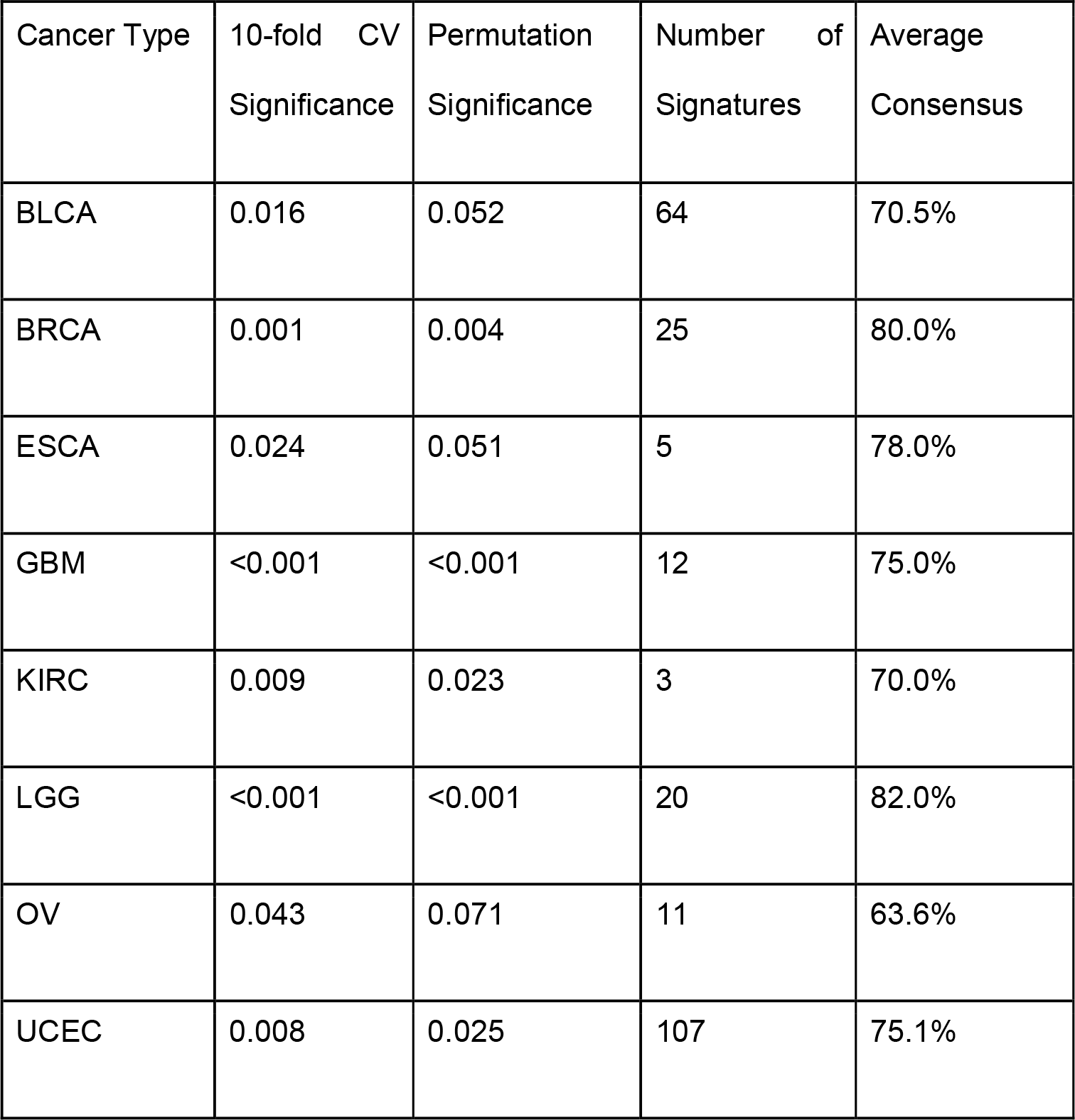
The results of cross-validation and permutation-based significance test for overfitting of prognostic signatures in eight validated cancer types. Number of prognostic signatures (including PIU, LU and NCU) with mutations occurring in at least three patients along with their average consensus in cross validation folds. Consensus for each prognostic unit is the number of times it being identified as prognostic unit in cross validation folds.

### Prognostic signatures associated with poorer survival outcomes are cancer-specific

The vast majority of Cox models with significant findings had positive regression coefficients (245 out of 247), suggesting deleterious impact of mapped somatic mutations on patient survival. **Figure 3A** illustrates that UCEC has the greatest number of prognostic signatures, followed by BLCA, BRCA, and LGG. In sum, PIUs outnumber LUs and NCUs. Prognostic PIUs uncovered by GPD consists of protein domains and PTM sites on genes implicated in various biological processes (**Figure 3B**). A large proportion of PIUs are enzymes catalyzing chemical reactions in cellular processes. For example, somatic mutation counts in the hydrolase domain in *NIT2* gene for ovarian cancer patients (OV) and those in FAH and *ATP13A3* genes for uterine cancer (UCEC) patients were significantly associated with overall survival. Mutations mapped to the peptidase domain in *FURIN*, *LGNM* and *CASP10* genes in UCEC patients, and in *TINAG* gene in breast cancer (BRCA) patients are also associated with poorer survival outcomes, highly likely by interfering with the catabolism of peptides into amino acids. Domains involved in altering the genetic material constitute a good proportion as well. For example, two histone domains in *H3F3A* and *H3F3B* have mutations seen in three lower grade glioma (LGG) patients individually; N-terminal Myc family in the transcription factor *MYC* gene harbours mutations in seven UCEC patients. Mpp10 domain in *MPHOSPH10* gene is related to the processing pre-rRNA to mature rRNA, and is a possible prognosis predictor in bladder cancer (BLCA) patients. Cadherin domains are responsible for the formation of adherens junctions to bind cells with one another. GPD revealed four cadherin-related domains in *PCDH11X* (LGG), *CHD9* (UCEC), *PCDHA7* (BLCA) and *FAT1* (BLCA) that are significantly associated with overall survival. Five sequence units encapsulating PTM sites were also discovered to be prognostic for overall survival. These include a tyrosine phosphorylation site in *IDH1* (LGG), a threonine phosphorylation site in *SLC9A4*, an arginine methylation site and a threonine phosphorylation site in *TP53* (UCEC), and lastly, a threonine phosphorylation site in *TP53* (BRCA).

**Figure 3.**
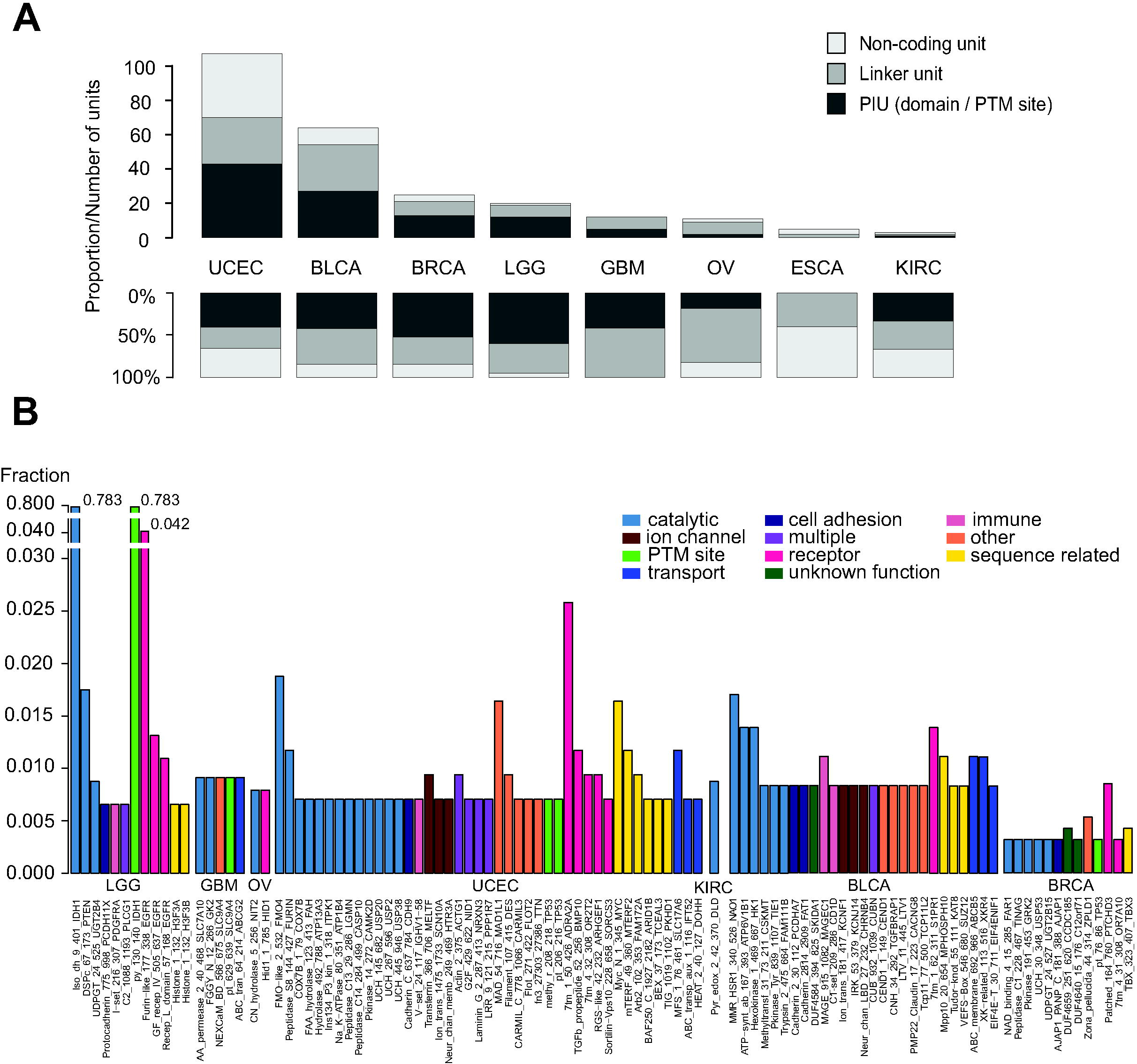
Association analysis between GPD somatic unit-level data and overall survival. (**A**) The total number and the proportion of significantly associated units (PIU, LU and NCU) for each cancer. (**B**) Significant PIUs across the seven cancers with prognostic signatures (no significant PIU identified in ESCA). Height of bars indicates the fraction of patients with one or more mutations on each PIU. Color of bars indicates the function each PIU is involved in.

It is well known that the isocitrate dehydrogenase *IDH1* is highly mutated in LGG patients (78.3%). An examination of the mutational landscape brings attention to a hotspot arginine 132 (R132), encapsulated the domain Iso_dh and a PTM site window for a tyrosine phosphorylation site (position 130-140) mentioned above. In most cases, the frequent missense mutation converts the arginine residue to histidine. A large body of evidence suggests that the detection of mutated IDH in glioma affords a survival advantage compared with wild-type IDH. (Lu and McDonald, 2018). GPD confirms this phenomenon with negative log coefficients (−1.64) from the Cox proportional hazards model for the unit counts in both the Iso_dh domain and the tyrosine phosphorylation site window. Epidermal growth factor receptor *EGFR* is also highly mutated in different sequence regions in LGG patients. 19 out of 457 patients have mutations in the Furin-like domain, 6 in GF_recep_IV domain and 5 in Recep_L domain. Overall, we observed that prognostic markers are highly specific to one cancer type, possibly suggesting that the most prognosis-related functional alterations caused by somatic mutations are cancer specific.

### GPD discovers associations overlooked by gene-level analysis

When somatic mutations are counted at the whole gene level, we obtained substantially different sets of survival associated genes (**Supplementary Figure 3**). In sum, > 75% of the genes (182 in 241) containing at least one significant sequence units were not discovered by gene-level analysis (**Supplementary Table 3**). The extreme case is in oesophageal carcinoma (ESCA), where two genes *ZNF459* and *RTN1* discovered by gene-level analysis were seen in unit-level results, and three other prognostic units bearing genes *EML4*, *LPAL2*, *REG3A* are novel findings. In glioma (LGG), *IDH1*, *EGFR*, *PTEN* and many other known genes were discovered in both levels, but histone domains in *H3F3B* and *H3F3A* were overlooked by the gene-level analysis. For the genes reported by both approaches, unit-level analysis highlights specific regions associated with survival outcomes. Moreover, unit-level analysis adds a total of 185 high-resolution discoveries in 182 novel genes to the set of prognostic markers.

### Survival-associated sequence units moderately overlap with driver mutations with prognostic values

We next compared survival-associated sequence units against known driver mutations and their corresponding genes (**Supplementary Tables 4** and **5**). To this end, we used the 299 cancer driver genes and ~3,400 driver mutations identified *in silico* by Bailey *et al.*, who employed a multitude of software tools to obtain the consensus list with data from the TCGA Pan-Cancer Atlas cohort (Bailey et al., 2018). We identified multiple PIUs and one LU within eight known driver genes, namely *FAT1, TBX3, TP53, IDH1, EGFR, PTEN, PLCG1,* and *NF1* (**Supplementary Figure 3**). Further, using the position information for the core set of 579 missense driver mutations, we mapped the mutations to prognostic sequence units. 32 missense driver mutations were found in seven prognostic units, including Furin-like domain, Receptor L-domain and GF_recep domain in *EGFR*, phosphorylation site T211 and methylation site R213 in *TP53*, Iso_dh domain and phosphorylation site Y135 in *IDH1*. This relatively scarce overlap between sequence units and driver genes/mutations is not surprising, since driver mutations which confer a selective growth advantage to tumors are not necessarily predictive of patient survival. Moreover, GPD only identified a small number of sequence units with highly stringent criteria for association with survival (passing the thresholds of *q*-value 0.05 and 10-fold cross-validation). However, the aforementioned genes and sequence units overlapping with GPD’s unit level analysis are structural components of genes that can prognosticate clinical endpoints in respective cancers, adding reliability to the reported discoveries by the GPD.

### Interactions between somatic and germline variants in the association with overall survival

We have shown that GPD allows for detection of sequence units enriched with variants that are associated with OS. Next, we exploited GPD’s capability to test for interactions between germline variants and somatic mutations within each gene in the association with overall survival. Such analysis is unique and of high clinical importance, as each subject’s germline genome may modulate the effects of somatic mutations.

We surveyed all sequence units in the eight cancer types, for which at least three patients have both somatic mutations in each sequence unit and germline variants in the gene containing the unit. This requirement rendered 8,240 PIUs, 9,641 LUs, and 5,209 NCUs in 8,821 genes eligible. For each sequence unit, we tested for the main effects of somatic mutation counts and germline variant counts as well as the interaction between the two, with adjustment of potential confounders (see **Materials and Methods)**. We did not adjust for multiple testing because the test could be performed for a relatively small number of units in each cancer cohort due to inclusion criteria explained above.

**Figure 4A** shows that bladder cancer (BLCA) had the largest number of interaction effects, followed by uterine (UCEC), ovarian (OV), brain (GBM) cancers. **Supplementary Table 6** provides the list of 520 significant interactions across the 8 cancer types. Interestingly, LUs appear to occupy the majority of significant interactions. In testing somatic mutations alone, more PIUs are significant (**Figure 3A**). Around half the interactions (280 out of 520) had positive coefficients, suggesting synergistic adverse effects of simultaneous occurrence of somatic mutations and germline variants on overall survival in these cases.

**Figure 4.**
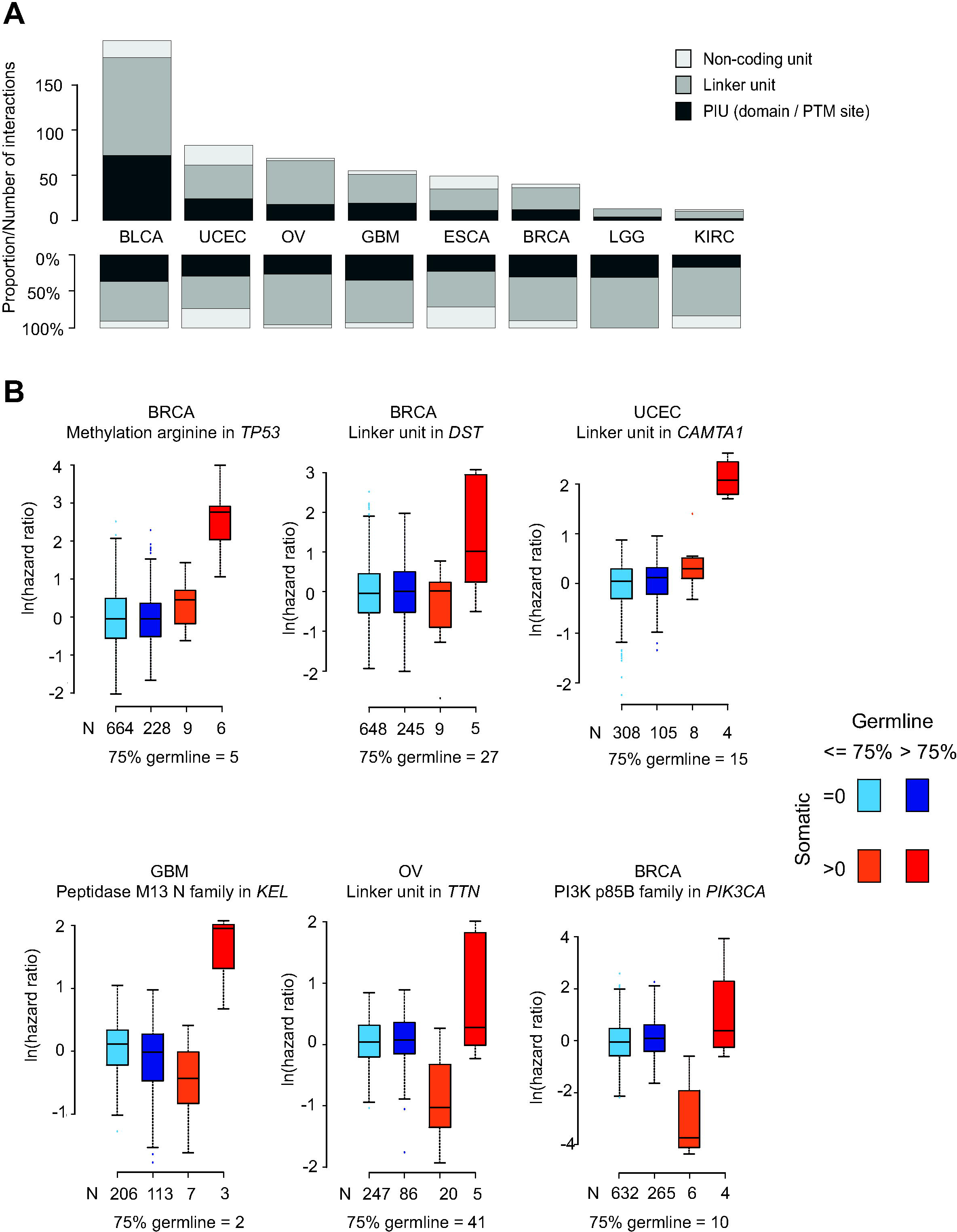
Testing the interaction effect of somatic and germline variants in cancer prognosis. (**A**) The number of statistically significant interaction effects in each cancer, among the eligible units (see **Materials and Methods**). (**B**) Comparison of the hazard ratios between patient groups with different somatic and germline variant levels. Patients are divided into four groups based on whether they have zero or at least one somatic mutation in the unit and whether they have fewer or more germline variants than the 75% quantile across all patients.

**Figure 4B** shows examples demonstrating how the unit-level frequency of germline variants modifies the hazard ratio of somatic mutation counts. For example, there were multiple occurrences of both germline variants and somatic mutations in the PIU surrounding the arginine methylation site R110 of the *TP53* gene. Fifteen out of 907 breast cancer (BRCA) patients had at least one somatic mutation in this sequence unit and 234 patients had more than ten germline variants on that gene. We found that the difference in the hazard ratios between a patient with somatic mutation(s) and a patient without one was much greater if the patients had more germline variants. Similar patterns were observed in the *DST* and *CAMTA1* genes in their respective cancer types. In comparison, as seen in the bottom panel of **Figure 4B**, having very few germline variants along with a somatic mutation decreased the hazard ratio and elevated the risk of death substantially in patients with more germline variants.

In sum, these findings suggest that substantial germline variation near the sequence units predisposes patients to a higher risk of death. Somatic mutations coupled with more germline variants in the same gene tend to have enhanced negative effects on survival, while somatic mutations surrounded by few germline variations may have protective effects.

## Discussion

We present a new approach, GPD, which summarizes sequence variation and mutations as frequency in small sequence units, providing two main advantages over alternative approaches. First, mutation count data for GPD’s sequence units mitigate the sparsity problem of locus-level exome sequencing data, addressing the inherent drawback that most exome variants do not occur at the same position with reasonably high frequency across different subjects. The sequence units therefore allow for immediate downstream statistical analysis to test their association with clinical outcomes. However, we acknowledge the limitation that our mapping scheme ignores potentially different protein sequence alterations caused by different types of mutations (e.g. treating missense mutations and early stop codon changes the same).

Second, GPD exploits the fact that variant data are not randomly distributed across a gene’s sequence but accumulates in specific regions. In contrast to conventional gene-level summaries, it does not sacrifice statistical power. At the same time, the sequence units associated with the OS inform on the direct functional consequences of the variants, facilitating the interpretation and enabling prioritization of targets for downstream experimental validation. Combined with survival analysis and association analysis for other clinical endpoints such as therapeutic response, the information can streamline patient-specific prognosis. To illustrate this, we have repeated the same analysis pipeline with PFI as the clinical outcome, where we report positive signatures validated in six cancer types. The results can be found in **Supplementary Information** document.

We first validated our findings through their reproducibility across a different large-scale dataset (MSK-IMPACT), confirmation by well-known cancer mutations, enrichment in protein domains implicated in cancer-related processes amongst survival associated genes, and overlap with an independently identified set of driver mutations. Furthermore, most sequence units with somatic mutations were highly patient-specific: they occurred in a small percentage of samples across cancers.

Another major advance in our work is the assessment of the combinatorial impact of different types of sequence variation, identifying interactions between existing genetic predisposition of a patient (germline variants) and tumorigenic events (somatic mutations) with respect to disease prognosis. To date, only few other studies have explored the interaction between germline and somatic variation, but in a limited fashion. For example, Carter *et al*. (Carter et al., 2017) analysed ~6,000 tumors and identified 412 genetic interactions between germline polymorphisms and major somatic events. However, the analysis was limited to 138 known oncogenes and tumor suppressor genes and focused on the effect of inherited polymorphisms in germline on the likelihood of somatic mutations, rather than discovering the prognostic value of the interaction between the two classes of sequence variation. Therefore, to the best of our knowledge, the interaction between genetic pre-disposition and somatic mutations has not been systematically addressed through *statistical association analysis* in the genome scale.

We found many genes in which patients with few germline variants and at least one somatic mutation in the same gene had better OS. By contrast, in other genes with many pre-existing germline variations, we found that additional somatic mutations almost exclusively had deleterious effects on overall survival. This finding implies that genetic predisposition (germline variants) affects the impact of subsequent mutation events on a gene and their indirect impact on patient survival -- highlighting a new avenue for individualized treatment based on the patient’s exome data. Given that GPD provides a highly flexible tool that can be applied to any definition of sequence units and large-scale datasets, integrative analyses of such interactions between germline and somatic variants can provide a new avenue for future individualized prognostic and therapeutic tools.

## Acknowledgments

This work was supported in part by grants from Singapore Ministry of Education (MOE2016-T2-1-001 and MOE2018-T2-2-058 to H.C.) and the National Institutes of Health (NIH 5R35GM127089-02 to C.V.). The results shown here are in whole or part based upon data generated by the TCGA Research Network: https://www.cancer.gov/tcga.

## Data Availability Statement

Data sharing is not applicable to this article as no new variant data were created in this study.

## Ethical Compliance

This study describes re-analysis of publicly available variant data and no new prospective or retrospective study was conducted.

