## Supplementary Information for "A protein-centric approach for exome variant aggregation enables sensitive association analysis with clinical outcomes"

**Survival analysis for PFI with GPD’s mutation/variant summary**

As illustration of GPD mapping with other clinical outcomes, we performed progression free interval (PFI) as the clinical endpoint in this analysis. In summary, GPD identified 169, 161, and 92 unique PIUs, LUs and NCUs that were associated with PFI in at least one of 22 cancer types, with mutation(s) in at least three patients (*q*-value 0.05). Also, Schöenfeld residuals suggested no violation of proportional hazards assumption in these units. We performed 10-fold cross validation on each cancer type (see **Materials and Methods**) to examine if the discovered prognostic units are capable of predicting survival outcomes in a reproducible manner. In six cancer types (ESCA, GBM, LGG, LUSC, PRAD, OV), the predictive indices cluster patients into high- and low-risk groups with significantly different survival outcomes (**Supplementary Figure 4** and **Supplementary** **Table 7**). Moreover, we conducted a permutation test to evaluate the degree of overfitting in the cross-validation process (see **Materials and Methods**). These six cancers survived the test with no sign of overfitting (**Supplementary** **Table 7** and **Supplementary Figure 5**). Therefore, our final prognostic signatures feature 60 PIUs, 44 LUs and 12 NCUs discovered in these cancer cohorts (**Supplementary Table 8 and Supplementary Figure 6**).

When somatic mutations are counted at the whole gene level, we obtained substantially different sets of survival associated genes (**Supplementary** **Figure 7**). > 57% of the genes (63 in 109) containing at least one significant sequence units (in total 65 units) were not discovered by gene-level analysis (**Supplementary Table 9**).

Comparing with possible driver genes identified by Bailey et al., we identified multiple PIUs and two LUs within seven known driver genes, namely *CIC, SAMRCA4, NOTCH1, IDH1, EGFR, PTEN,* and *NF1* (**Supplementary** **Figure 7 and Supplementary Table** **10**). Further, using the position information for the core set of 579 missense driver mutations, we mapped the mutations to prognostic sequence units. 32 missense driver mutations were found in seven prognostic units for PFI (**Supplementary Table** **11**).

We also repeated the analysis of interaction between somatic mutations and germline variants. We discovered 522 significant interactions across the 6 cancer types (**Supplementary Table 12**). **Supplementary** **Figure 8A** shows that lung cancer (LUSC) had the largest number of interaction effects, followed by brain (GBM) and ovarian (OV) cancers. **Supplementary** **Figure 8B** shows four representative examples of interaction effects.

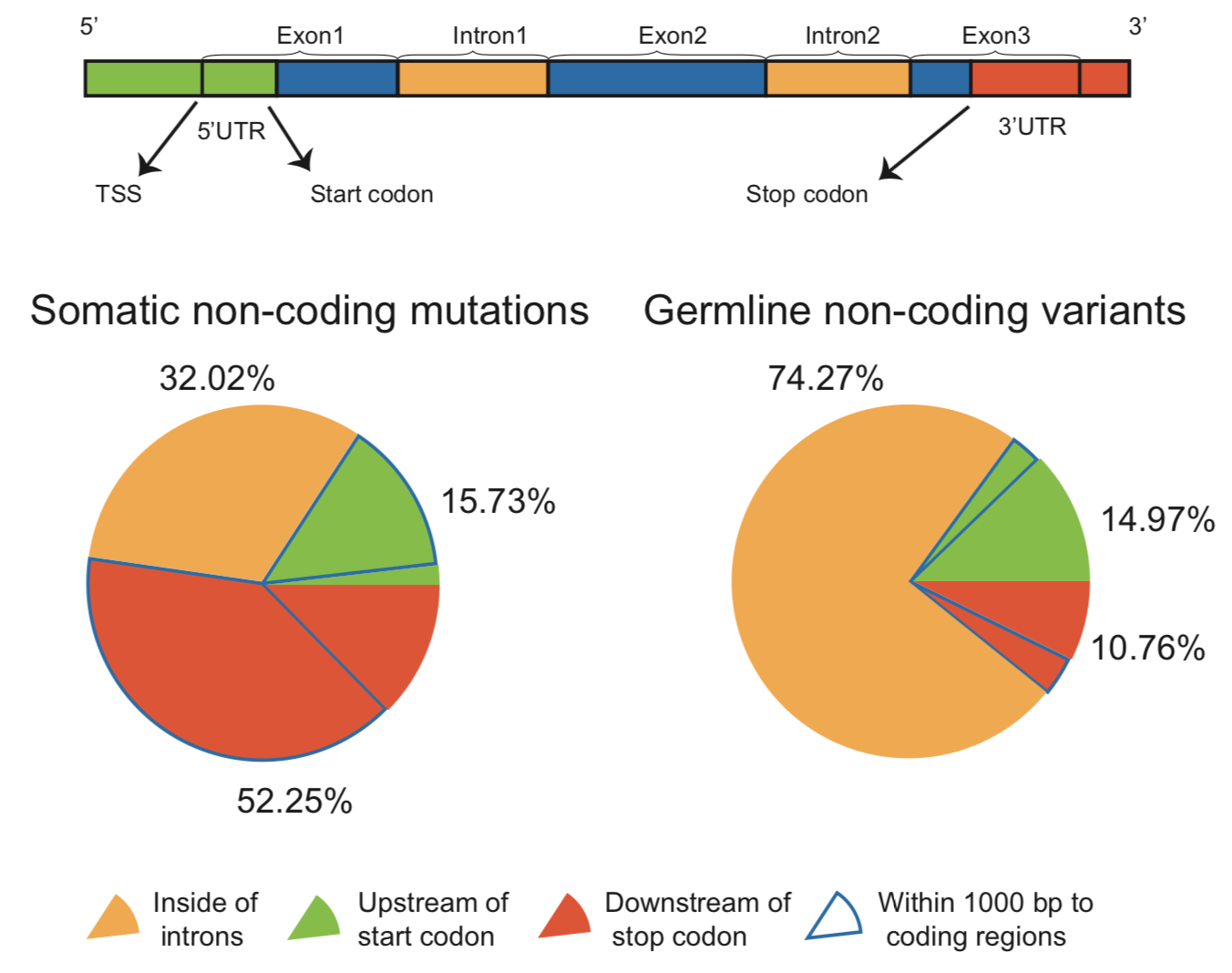

**Supplementary Figure 1.** Relative position from mutations/variants mapped to non-coding regions (yellow, green and red) to coding exons (blue). Pie charts showing the percentage of such mutations/variants distributed in introns (yellow), upstream of start codon (green), and downstream of stop codon (red). Sections outlined in blue indicates the fraction of mutations/variants distributed within 1000 base pairs to coding regions and upstream of start codon or downstream of stop codon.

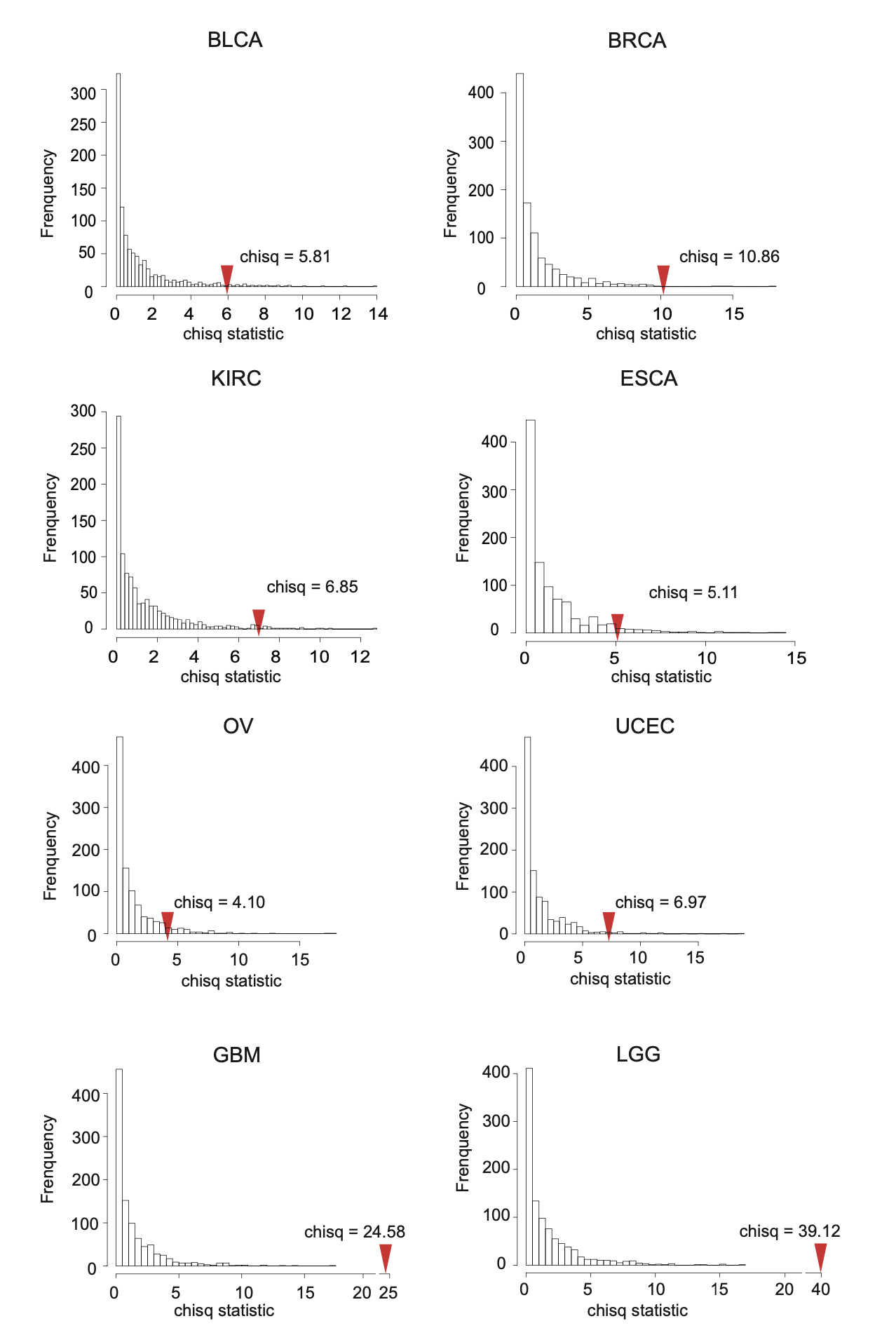

**Supplementary Figure 2.** Chi-square statistics derived from 1,000 permutations of 10-fold cross-validation, where survival outcomes are randomly shuffled. Red spike indicates chi-square statistic of cross-validation result derived from the analysis with un-permuted, original survival outcomes.

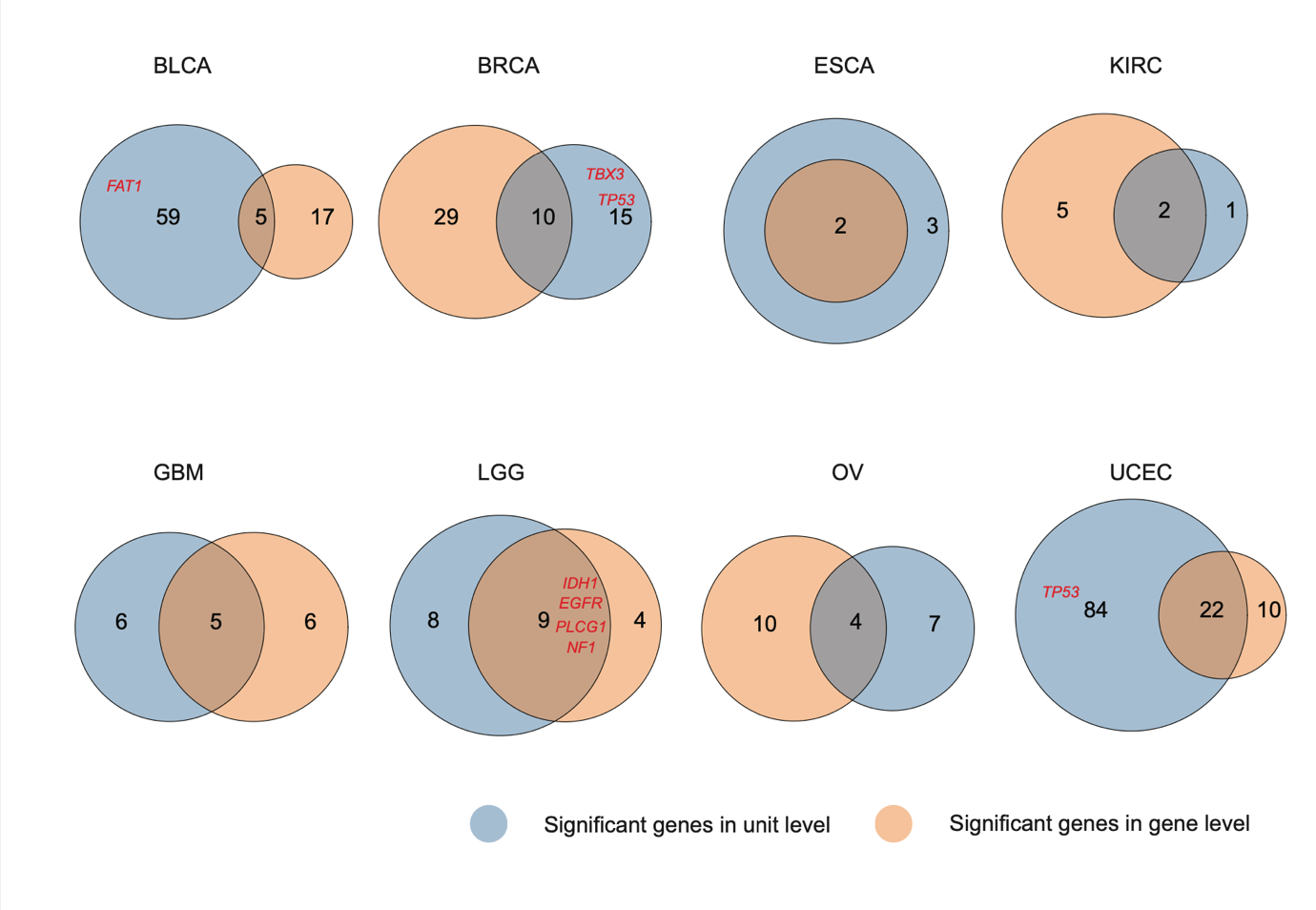

**Supplementary Figure 3.** Overlap between significant genes discovered by GPD gene-level analysis (orange) and genes harboring significant units discovered by GPD unit-level analysis (blue). Genes in red are ones harboring significant units discovered by GPD and identified as driver genes with TCGA Pan-cancer software.

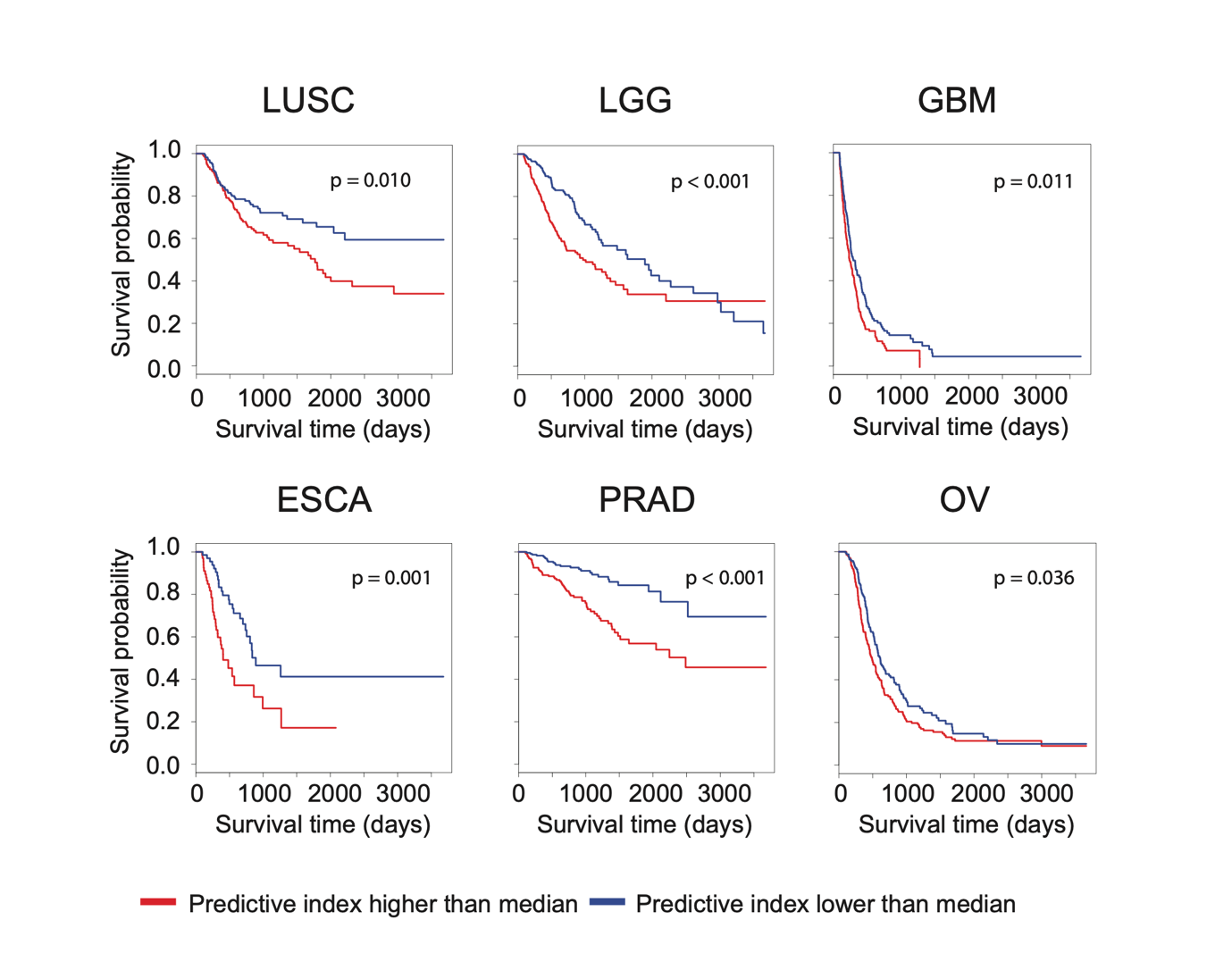

**Supplementary Figure 4.** Kaplan-Meier plot for six cancer types with prognostic signature for PFI. Predictive index (PI) was derived for each sample in the test group in each fold of the 10-fold cross validation. Patients who have PIs greater than the median in training group are clustered in the high-risk group (red), those with lower than median PIs are in the low-risk group (blue). P-values are derived from log-rank tests.

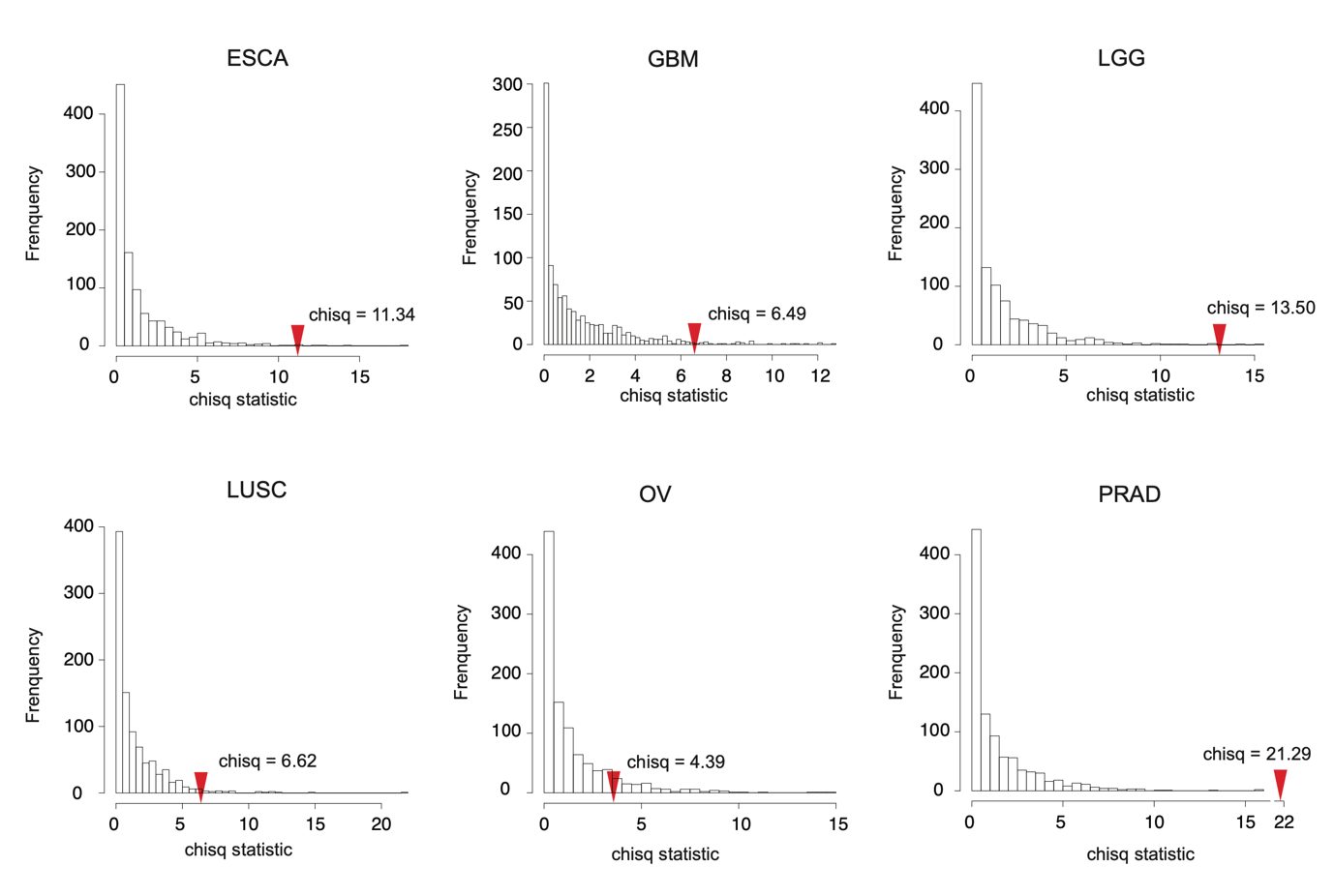

**Supplementary Figure 5.** Chi-square statistics derived from 1,000 permutations of 10-fold cross validation, where the survival outcomes (PFI) are randomly shuffled. Red spike indicates the chi-square statistic of cross validation result derived from the original survival.

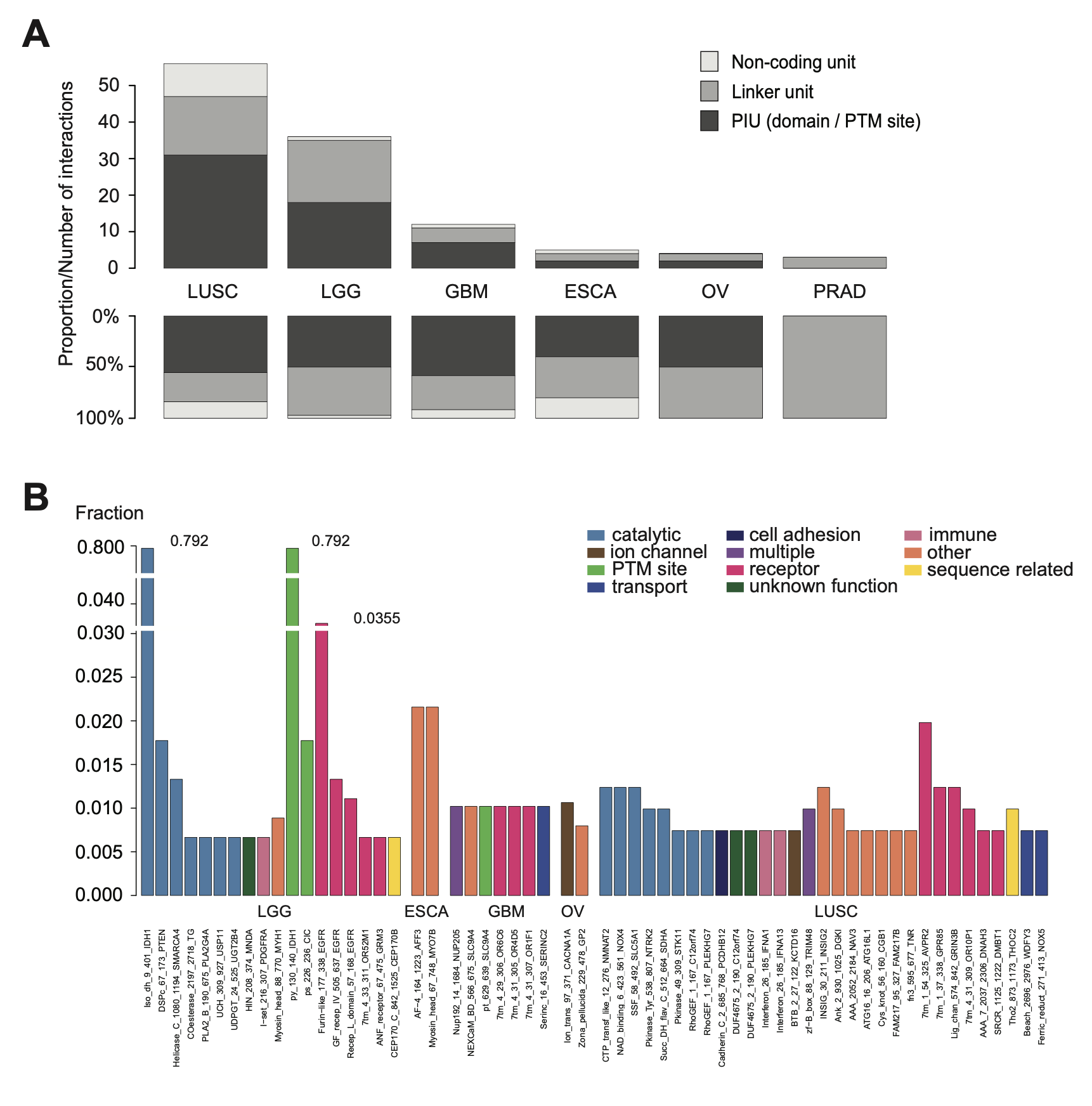
**Supplementary Figure 6.** Association analysis between GPD sequence unit-level somatic mutation counts and PFI. (**A**) The total number and the proportion of significantly associated sequence units (PIU, LU and NCU) for each cancer. (**B**) Significant PIUs across seven cancers (no significant PIU identified in PRAD). Height of bars indicates the proportion of patients with one or more mutations on each PIU. Color of bars indicates the biological processes PIUs are involved in.

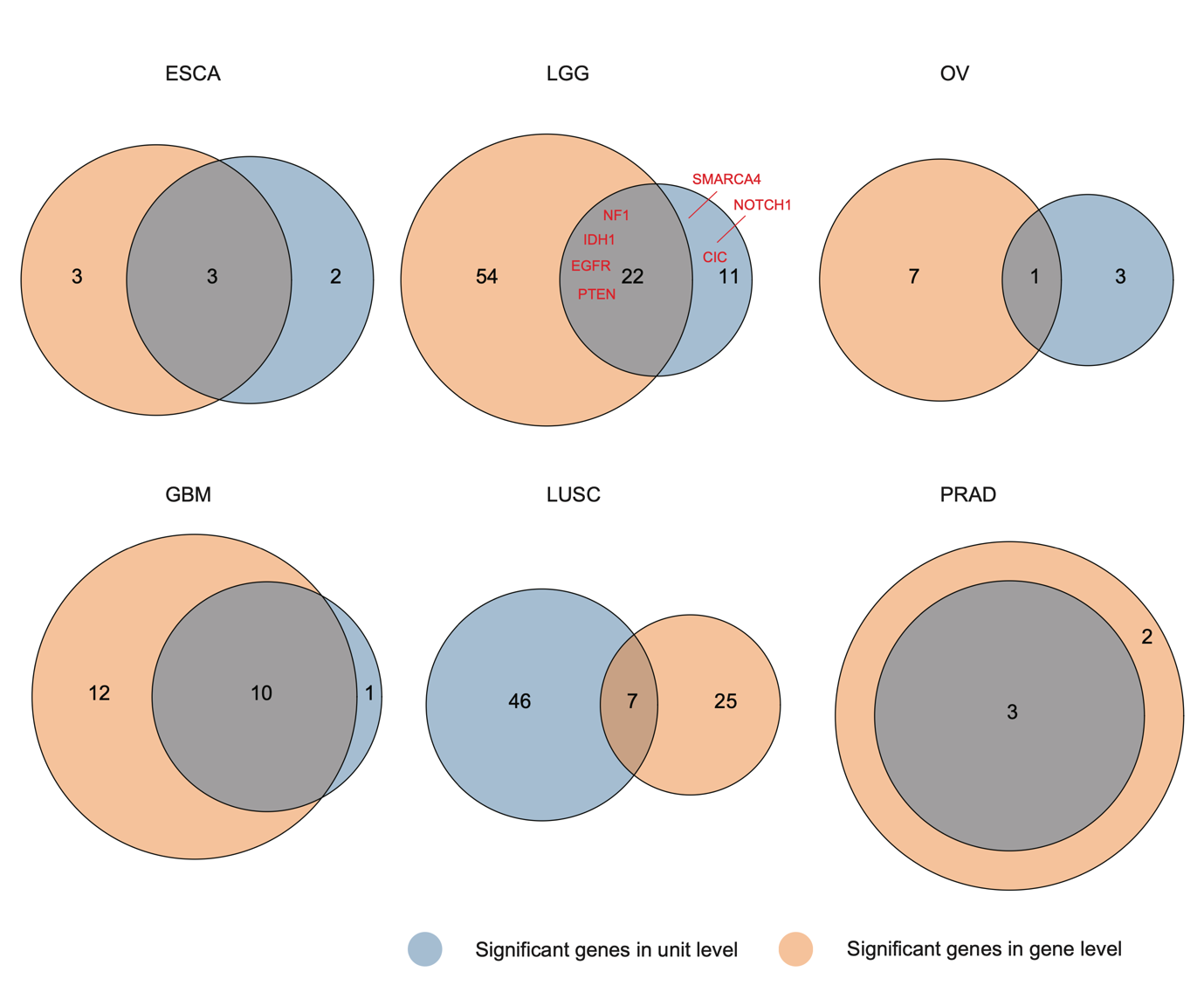

**Supplementary Figure 7.** Overlap between significant genes discovered by gene-level analysis (orange) and genes harboring significant units discovered by unit-level analysis (blue). Genes in red are ones harboring significant units discovered by GPD and identified as driver genes with TCGA Pan-cancer software.

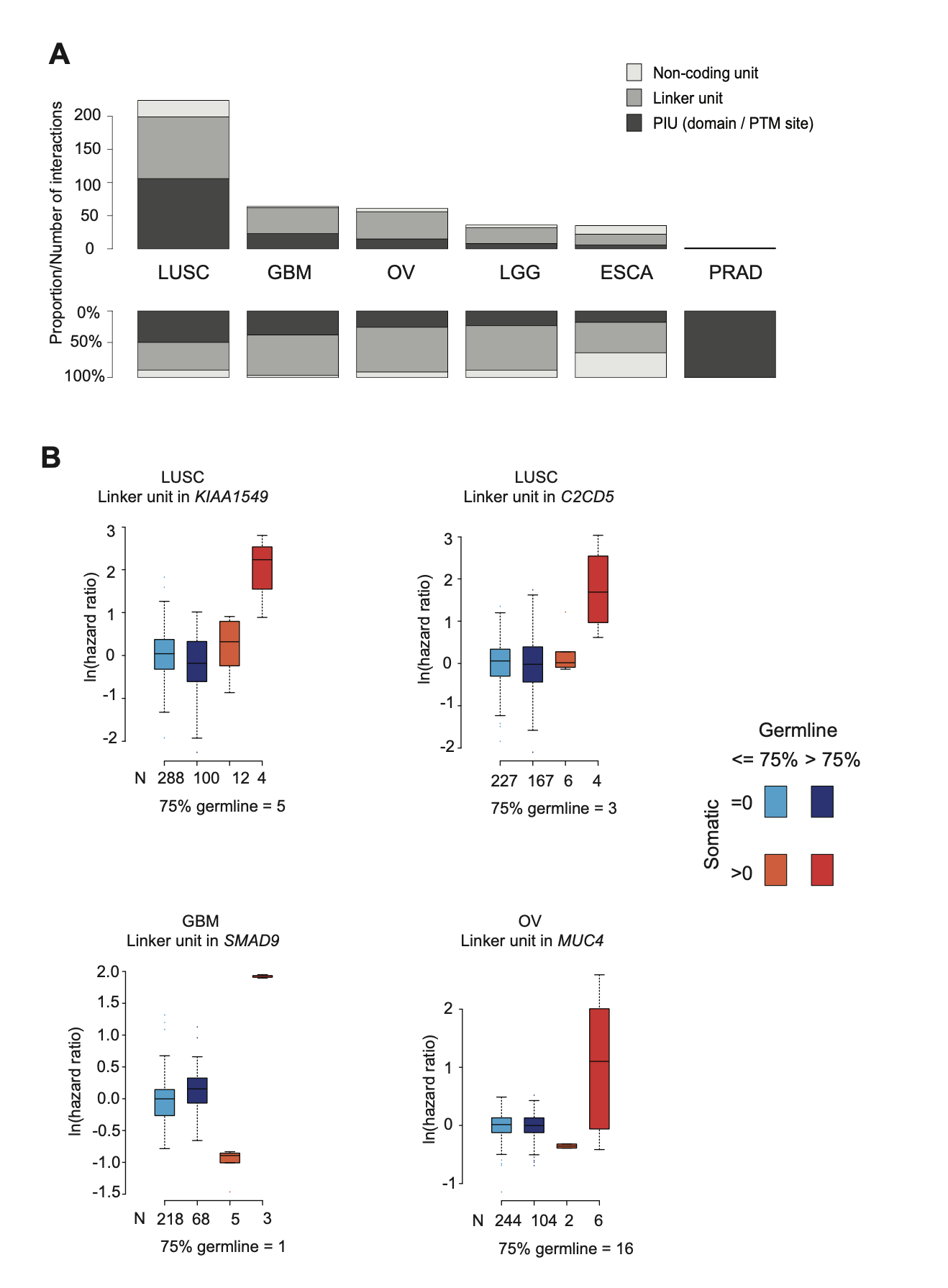

**Supplementary Figure 8.** Test of the interaction effect between somatic and germline variants in cancer prognosis (PFI). (**A**) The number of statistically significant interaction effects in each cancer, among the eligible units (see **Materials and Methods**). (**B**) Comparison of the hazard ratios between patient groups with different somatic and germline variant levels. Patients are divided into four groups based on whether they have zero or at least one somatic mutation in the unit and whether they have fewer or more germline variants than the 75% quantile across all patients.
